# Real-time respiration changes as a viability indicator for rapid antibiotic susceptibility testing in a microfluidic chamber array

**DOI:** 10.1101/2021.01.02.425088

**Authors:** Petra Jusková, Steven Schmitt, André Kling, Darius G. Rackus, Martin Held, Adrian Egli, Petra S. Dittrich

**Author notes:** **Corresponding Author**, Prof. Dr. Petra Dittrich.

## Abstract

Rapid identification of a pathogen and the measurement of its antibiotic susceptibility are key elements in the diagnostic process of bacterial infections. Microfluidic technologies offer great control over handling and manipulation of low sample volumes with the possibility to study microbial cultures on the single-cell level. Downscaling the dimensions of cultivation systems directly results in a lower number of bacteria required for antibiotic susceptibility testing (AST) and thus in a reduction of the time to result. The developed platform presented in this work allows the reading of pathogen resistance profiles within 2-3 hours based on the changes of the dissolved oxygen levels during bacterial cultivation. The platform contains hundreds of individual growth chambers prefilled with a hydrogel containing oxygen-sensing nanoprobes and different concentrations of antibiotic compounds. The performance of the microfluidic platform is tested using quality control *Escherichia coli* strains (ATCC 25922 and ATCC 35218) in response to different clinically relevant antibiotics. The achieved results are in agreement with values given in clinical reference guides and independent measurements using a clinical AST protocol. Finally, the platform is successfully used for AST of an *E. coli* clinical isolate obtained from a patient blood culture.

## INTRODUCTION

Infectious diseases are a significant cause of morbidity and mortality worldwide.^1^ With the emergence of antibiotic resistance,^2^ bacterial infections are becoming an important threat to global health.^3^ Particularly in the case of blood stream infections, which are often associated with severe sepsis or septic shock, early administration of effective antibiotics is extremely time critical for successful treatment.^4^ Rapid antibiotic selection and proper dosage are therefore extremely important.

Standard growth-based assays for antibiotic susceptibility testing (AST) include manual methods such as broth microdilution on a multi-well plate and gradient methods on gel-dishes, or automated methods using commercial instruments (e.g. VITEK 2, bioMérieux SA). Regardless, these methods require between 10^4^ - 10^8^ CFU mL^-1^, leading to long pre-incubation times which could result in several days until the AST can be performed and results obtained.^5^ Molecular AST techniques, mostly relying on polymerase chain reaction, are rapid and sensitive, but only suitable for already well-characterized resistance genes. Moreover, they are relatively expensive and require well-trained personnel.^6,7^ Therefore, the development of new rapid and sensitive diagnostic tools is of great importance.^8,9^

Strategies for reducing the AST time include the miniaturization of test systems and improvements in readout. Microfluidic devices typically operate with volumes of μL to fL and are therefore promising systems for antibiotic susceptibility testing.^10,11^ Fabrication of wells or the creation of water-in-oil emulsions on a microfluidic platform decreases the initial number of bacteria required to just one or few cells per compartment.^12^ This also significantly shortens the time required for pre-culturing and further analysis. Additionally, microfluidic techniques can be combined with molecular ASTs^13,14^ facilitating complementary bacterial identification or analysis.^15^

Droplet systems offer compartmentalization of the bacterial samples, continuously and in a very high throughput, enabling analysis of thousands of samples^16^ and on the single-cell level.^17^ However, the pump systems typically required to generate emulsions are difficult to integrate with the point of care systems, while the alternative, gravitational forming operates at a significantly lower throughput.^18^ In addition, droplet microfluidics requires inert oils and detergents to stabilize the droplets, yet shrinking of droplets and leakage of compounds (e.g., via micelle formation) is often observed.^19^

Alternatively, devices with integrated cultivation chambers or wells have been introduced.^20–23^ Here, the chamber number is fixed, given by the initial design of the device. Isolation, i.e., closing the chamber, is achieved by valves^24,25^ or again, by supply of non-water miscible fluids (oils).^15,26–28^ SlipChip technology represents another approach to isolate individual cultivation compartments. Such devices typically contain two parallel plates which can be moved to fill and close micro to picolitre-volume wells.^14,29^ While the throughput is limited, these designs have other advantages, e.g. the chambers can be supplied with compounds, medium or antibiotics at a later time in the experiment, hence providing the possibility of tracking the bacterial growth in response to changes of cultivation conditions over time. Other designs use partially open chambers or channels without valves, designed to trap and immobilize bacterial cells for high-resolution single-cell imaging.^30–33^,

Many of the above mentioned devices are excellent research tools, but practical drawbacks limit their use in clinical settings. In addition, many microfluidic devices made of the polymer polydimethylsiloxane (PDMS) suffer from several limitations such as instable surface properties, absorption of hydrophobic compounds into the polymer, diffusion of water through the polymer and the limited possibility for mass production.^19,34,35^

In this study, we introduce a microfluidic device with embedded chambers that is made of cyclic olefin copolymer (COC) by thermal imprinting. The device is easy to operate as the final use requires only pipetting and sealing of the device, making its production and assay protocol easily scalable and adaptable for low-cost mass-production and potential clinical use. Since COC is gas-tight material, it further enabled us to improve the readout to differentiate susceptibility and resistance of aerobic pathogens. Here, we embedded oxygen-sensing nanoprobes in the chambers as the means for reading cell metabolic activity and hence, viability via relative changes in luminesce of the nanoprobes.

Several approaches have been introduced before that translate bacterial metabolic activity into a measurable optical signal. For example, adenosine triphosphate can be determined in a bioluminescence assay^36^ or pH-dependent color changes of a chromophore^23,24^ were successfully used for the susceptibility testing. The conversion of resazurin by the bacteria to resorufin is a fluorescence assay widely employed in microfluidic AST systems.^26–28^ The assay is very sensitive, but requires a high concentration of the reagent to generate signal during the long-time measurements. Considering that resorufin is also susceptible to photobleaching, the assay is more suitable for the end-point measurements rather than for dynamic changes of cell viability.^37^ Further, it is necessary to test cross-reactivity of the tested antibiotics with the reagent.^38^ In contrast, the luminescence signal generated by oxygen-sensing nanoprobes is reversible and provide an almost real-time indication of changes in the dissolved oxygen level in the surrounding environment.^39^ It was previously shown that these oxygen sensing nanoprobes are not compromising cell viability and are compatible with an oxygen measurements in shake flasks and microtitration plates.^40,41^

In the following, we show possibility to use oxygen sensing nanoprobes to assess bacterial viability in the microfluidic AST platform. The characterization of the platform is performed using two quality control strains of *Escherichia coli*, ATCC 25922 and ATCC 35218. Both strains constitutively produced a superfolder variant of the green fluorescent protein (sfGFP) for simplified monitoring of bacterial growth and observation of morphological changes in response to antibiotic exposure. The platform was finally validated using a clinical isolate of *E. coli* strain obtained from a positive blood culture of a septic patient.

## RESULTS AND DISCUSSION

A conceptual sketch of the AST devise is presented in Figure 1. The device comprises four sets of nanoliter-sized chamber arrays. These are pre-filled with agarose gel containing oxygen sensing nanoprobes and different concentration of the selected, clinically relevant antibiotics. The bacterial sample is deposited on a standard glass slide and covered by the pre-filled chamber array. The measurements are performed on an automated microscope, collecting luminescence signals from the individual chambers.

**Figure 1.**
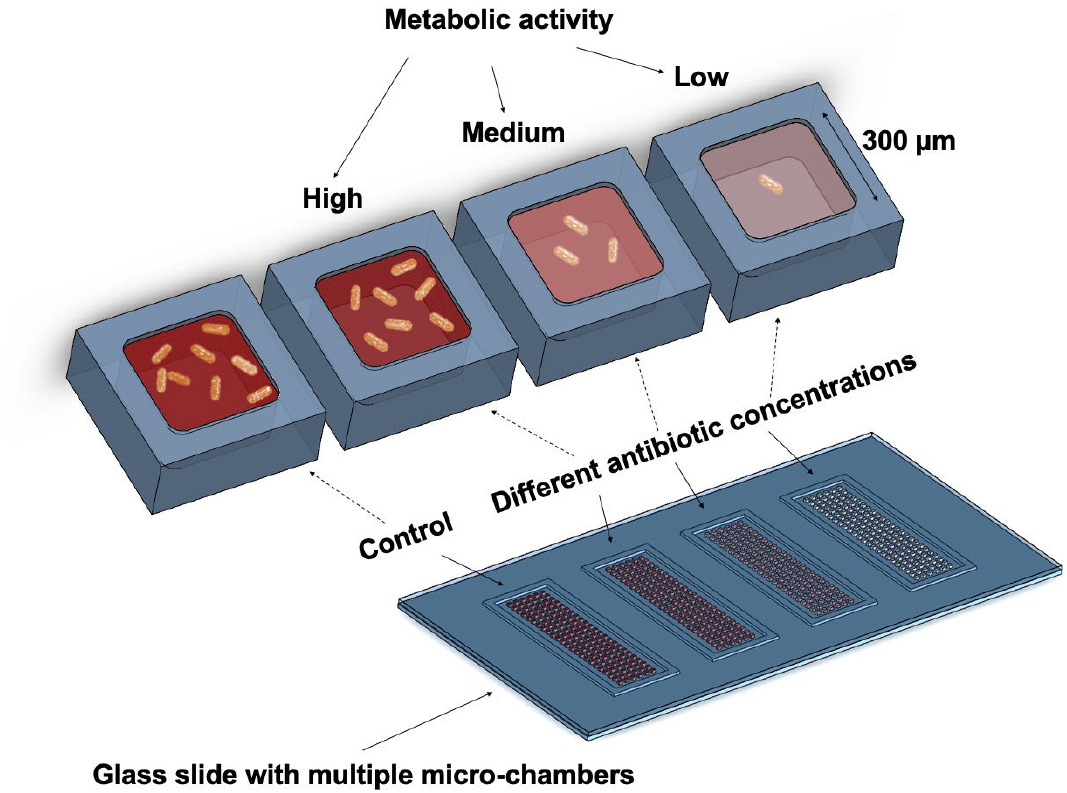
Conceptual sketch of the microfluidic system for antibiotic susceptibility testing. The platform consists of four sets of chamber arrays with hundreds of growth chambers (300 × 300 × 75 μm) fabricated using the air-tight material, cyclic olefin copolymer. The chambers are filled with oxygen sensing nanoprobes and different concentrations of the studied antibiotics embedded in the agarose gel. Changes in metabolic activity of the bacteria result in changes of the luminesce signal of the nanoprobes, enabling the determination of susceptibility or resistance of the studied pathogens against the various antibiotic compounds.

The dissolved oxygen acts as a quencher of the nanoprobe luminescence (Figure S1). The chambers containing metabolically active bacteria consume oxygen and thus provide greater luminescence signal compared to the chambers containing bacterial cells with metabolism affected by the antibiotics.

The AST workflow (Figure 2A) of the presented microfluidic platform starts by filling the individual chambers of the array with an agarose gel (2.5 % w/v) containing oxygen sensing nanoprobes and antibiotics diluted in the culture medium (Mueller Hinton Broth II, cation adjusted). Using four arrays on the same device allows for four different conditions to be tested simultaneously. Each individual array consists of 144 chambers (300 × 300 μm with a depth of 75 μm) and is surrounded by a trench to drain excess gel and bacterial culture once the device is assembled. Further, this prevents cross-contamination between the arrays. A droplet (~15 μL) of the liquid gel, maintained at 47 °C is pipetted on the COC plate and spread over the array using a thin PDMS slab. We selected an ultra-low gelling temperature agarose to avoid high operating temperatures and risk of degrading the antibiotic compounds; due to the low volume of the chambers (~7 nL), the gel solidifies nearly immediately after its deposition. The gel matrix helps to reduce evaporation of the media during the experiments, simplifies manipulation with the filled COC plate and also keeps the oxygen sensing particles in fixed positions during the measurement, avoiding their sedimentation. Detailed figures of the COC chambers are found in Figure S2.

**Figure 2.**
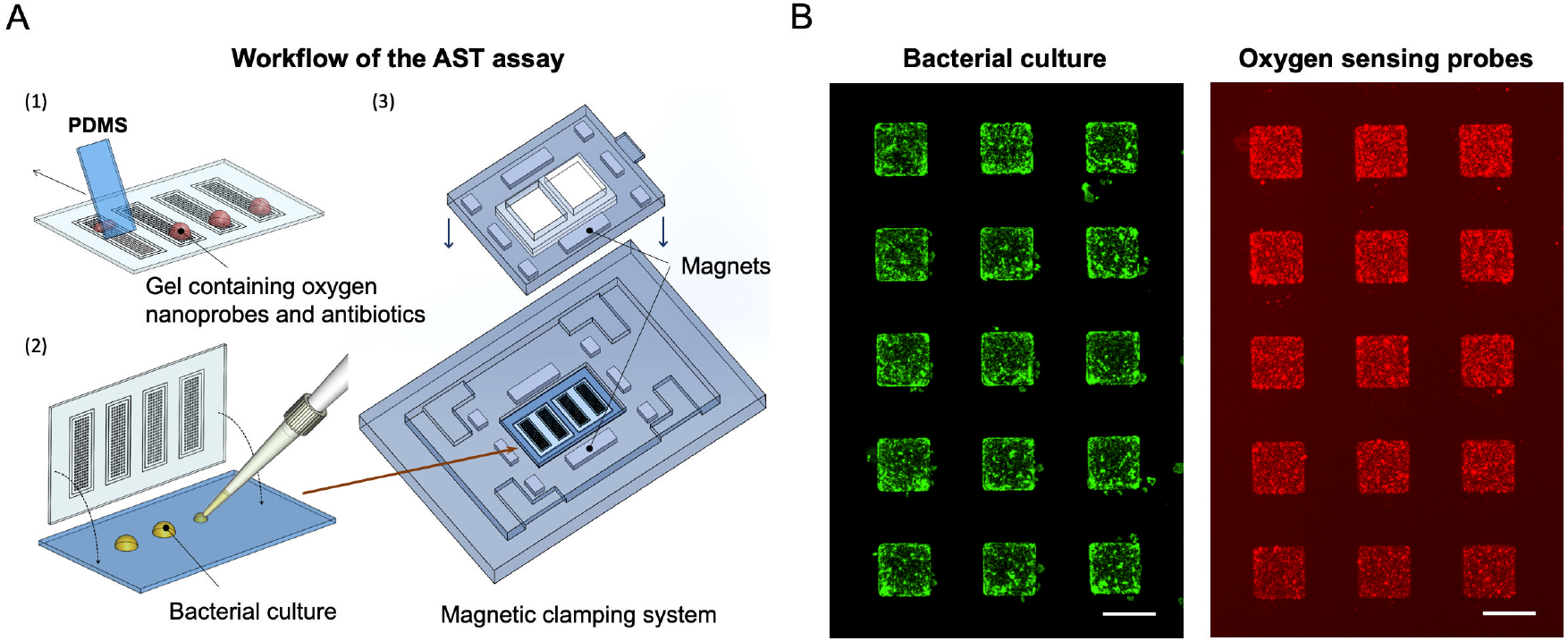
A: The experimental workflow. First, using a thin slab of PDMS, a small droplet of the liquid agarose (~ 15 μl) containing oxygen sensing probes and studied antibiotics is spread across the individual arrays. Next, 20 μL of a bacterial suspension (OD_600_=0.02) is deposited on a microscopy glass slide and overlaid with the microfluidic chamber arrays. Subsequently, arrays are transferred into a magnetic clamping device where they are kept in contact during the AST measurement. Oxygen consumption during the bacterial growth is then monitored using the automated microscope via the changes of the luminescence signal collected from the individual chambers. B: (Left) Detailed micrograph of the array with the sfGFP producing bacterial strain, growing predominantly in the chamber areas. (Right) The corresponding micrograph of the oxygen sensing nanoprobes embedded in these chambers. Scale bars: 300 μm.

In the following step, the bacterial suspension (OD_600_=0.02) is pipetted on a microscopy slide (20 μL split into four 5 μL droplets) and overlaid with the microfluidic chamber arrays. The mean cell number per array after the inoculation is between 10-45 CFU per chamber (data not shown). The glass slide and chamber arrays are maintained in contact using a simple magnetic clamping system which also serves as a holder for automated microscopy imaging (Figure S3). The bacterial cells entrapped in the chambers have sufficient medium and nutrients to grow for multiple generations.

Cells without access to the chambers have significantly lower nutrient supply and therefore, after five hours of cultivation we observed only a negligible number of cells in the regions outside of the chambers (Figure 2B). Oxygen consumption during bacterial growth is monitored as an increase in the luminescence signal collected from the oxygen sensing nanoprobes inside the individual chambers. Similarly, as for the bacterial cells, there is a very low portion of the nanoprobes in the areas between the chambers. As seen in Figure 2B, the majority of the probes are located within the chambers allowing us to record real-time changes in the luminesce level during the bacterial culture in each chamber. An example of the four chambers at the beginning of the culture and after 5 hours of cultivation can be seen in Figure S4. By comparing respiration changes of the cultures containing different antibiotics and a control without antibiotic supplement, we are able to distinguish between resistant and susceptible bacterial strains.

To characterize the system, we first performed AST using the fully susceptible quality control *E. coli* strain ATCC 25922. The strain was first tested against three different concentrations of ciprofloxacin (Figure 3A). The minimal inhibitory concentration (MIC) value for the *E. coli* ATCC 25922 against ciprofloxacin, listed in the EUCAST table,^42^ was used as a reference for the antibiotic concentration, MIC_ref_= 0.008 μg mL^-1^. Based on the luminescence signal originating from the oxygen sensing probes, the oxygen consumption was reduced by approximately half at the MIC_ref_ value, remained high at doses below the MIC_ref_ (0.25 MIC_ref_) and dropped close to zero when ciprofloxacin was added in excess (4 MIC_ref_), indicating a loss in bacterial viability. Moreover, we could clearly distinguish the signal difference in all tested conditions within the first 2.5 hours of cultivation.

**Figure 3.**
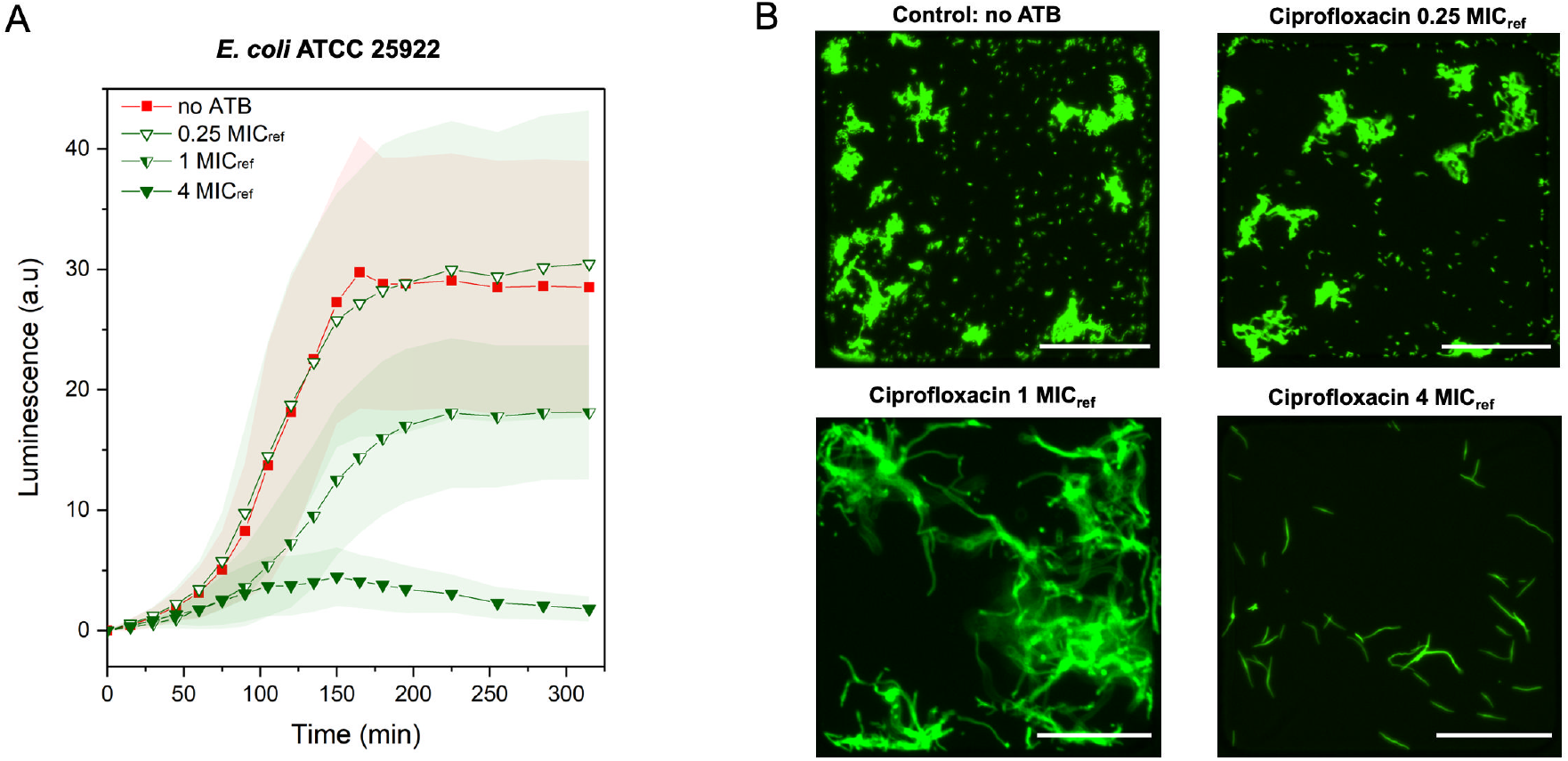
Characterization of the AST platform using quality control strain. A: Kinetic measurements of cell viability via oxygen sensing nanoprobes immobilized in the agarose gel matrix. Comparison between control without antibiotic (no ATB) supplement *E. coli* ATCC 25922 culture (RED, n=15), and cultivation in the presence of ciprofloxacin at three different concentrations, 0.25 MIC_ref_, 1 MIC_ref_, 4 MIC_ref_ (GREEN; n=15, 24, 66 and 42 respectively). B: Figures of the individual chambers show the bacterial morphology at the end of the measurement for each concentration of ciprofloxacin, MIC_ref_= 0.008 μg/ml. Scale bars: 100 μm. Temperature 37 °C; MHB II, cation adjusted media and the initial OD600 of 0.02 were maintained in all experiments. Error bars represent the standard deviations.

The obtained results were further supported by the observed changes in morphology (Figure 3B). Within the chambers containing antibiotics supplied at sub-MIC_ref_ doses bacterial cells formed micro-colonies surrounded by a high number of planktonic cells. Once the antibiotic concentration was supplied at the MIC_ref_ concentration, bacteria continued to grow, but we did not observe any cell division, leading to elongated bacterial cells. An excess of ciprofloxacin (4 MIC_ref_) resulted in the declined bacterial growth after about two hours of cultivation. The results of the performed AST assay confirmed that the MIC value obtained for ciprofloxacin using the presented platform is in a similar range as given by the EUCAST,^42^ and a value obtained using an independent broth dilution AST (SI, Table 1).

We further tested the strain susceptibility by exposure to excess meropenem and amoxicillin corresponding to the 4 MIC_ref_ concentration (for *E. coli* ATCC 25922), 0.016 μg mL^-1^ and 4 μg mL^-1^ respectively (Figure 4A). In comparison to the bacterial culture without the antibiotics, meropenem and amoxicillin presence in the chambers lead to significant decrease in the measured luminescence signal, confirming the susceptibility of the tested strain to these antibiotics. Images showing the morphological changes for one selected chamber per each tested condition can be found in the supplementary information (Figures S5, S6, S7, S8 and S9).

**Figure 4.**
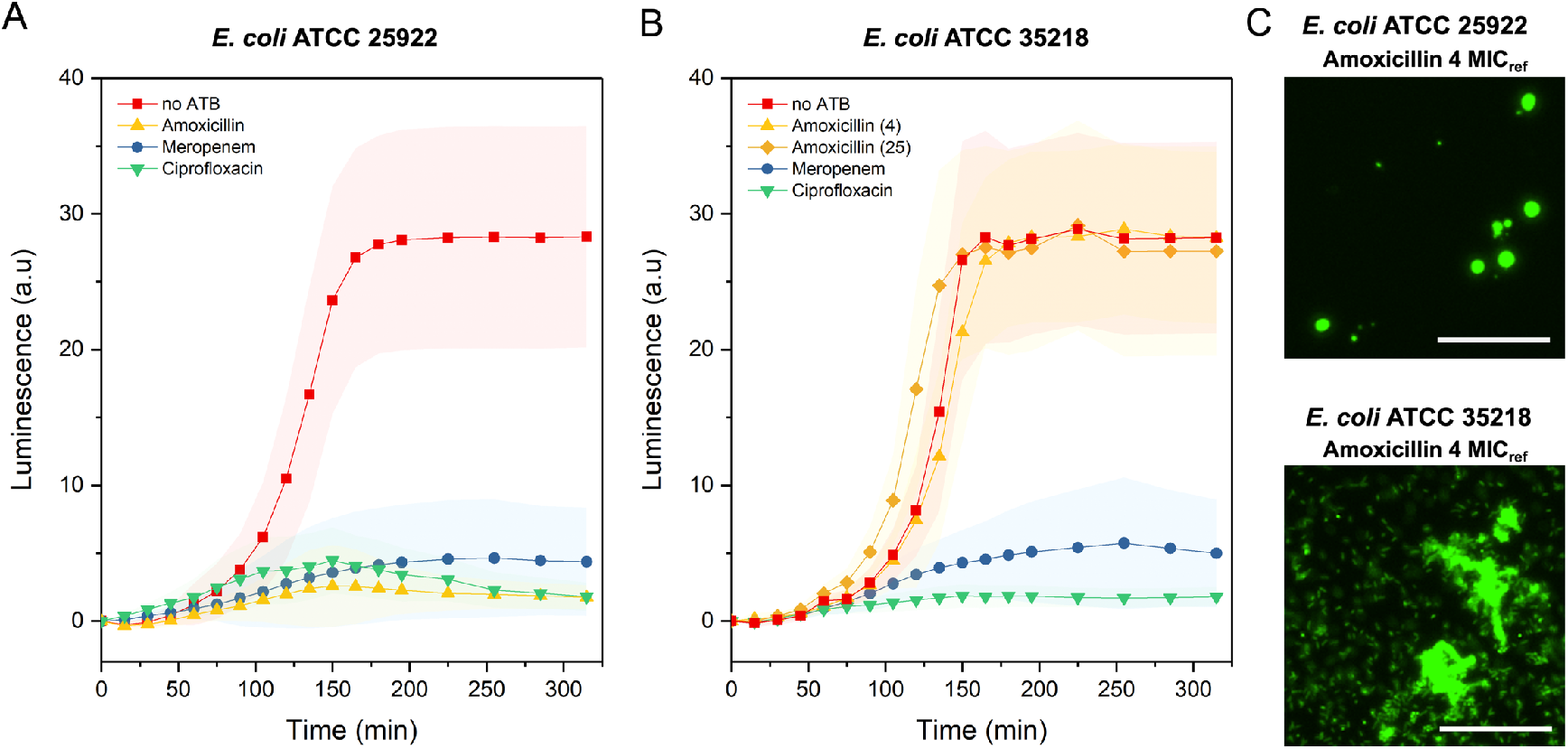
Validation of the AST platform using quality control strains. A: Comparison between control (no ATB) *E. coli* ATCC 25922 culture (RED, n=56) and cultivation in the presence of amoxicillin (YELLOW, n=71), meropenem (BLUE, n=76) and ciprofloxacin (GREEN, n=42) at a concentration four times greater than the MIC_ref_ (4 μg mL^-1^ for amoxicillin, 0.016 μg mL^-1^ for meropenem and 0.008 μg mL^-1^ for ciprofloxacin) B: Comparison between control (no ATB) *E. coli* ATCC 35218 culture (RED, n=74) and cultivation in the presence of meropenem (BLUE, n=48), amoxicillin (YELLOW, (4), n=43 and ciprofloxacin (GREEN, n=36) at a concentration four time greater than the MIC_ref_ (0.016 μg mL^-1^ for meropenem, 4 μg mL^-1^ for amoxicillin and 0.008 μg mL^-1^ for ciprofloxacin) together with amoxicillin (YELLOW, (25), n=55, supplied at a concentration 25 times greater than the MIC_ref_. C: Detailed micrographs from selected chambers after 225 min of cultivation showing the morphological difference between the susceptible (ATCC 25922) and resistant (ATCC 35218) *E. coli* strain in the presence of amoxicillin supplied at a concentration four times greater than the MIC_ref_ for amoxicillin. Scale bars: 50 μm. Temperature 37 °C; MHB II, cation adjusted media and the initial OD600 of 0.02 were maintained in all experiments. Error bars represent the standard deviations.

To further confirm the ability of the assay to correctly identify the bacterial resistance, we performed an AST assay using a TEM-1 beta-lactamase-producing *E. coli* strain (ATCC 35218) which is resistant to beta-lactam antibiotics such as amoxicillin. The results of the assay are presented in the Figure 4B; the tested antibiotics as well as their reference MICs were identical to the previous AST with *E. coli* ATCC 25922. Compared to the control culture without antibiotics, bacterial cultures supplied with meropenem and ciprofloxacin showed low values of the measured luminescence (oxygen consumption), confirming the susceptibility of the strain toward these antibiotics. Contrary, for the bacterial culture in the presence of amoxicillin, the respiration profile was in agreement with the culture without antibiotics. We observed a sharp increase in the luminescence shortly after the start of cultivation, which reached and maintained a maximum value at about 2.5 hours. A similar profile was observed when the concentration of amoxicillin was increased to 25 times greater than the MIC_ref_. Based on these AST measurements, we could confirm that the ATCC 35218 *E. coli* strain is indeed resistant to amoxicillin. Additionally, the selected antibiotic concentration, corresponding to four times the MIC_ref_ value, is sufficient to observe a difference in respiration profiles necessary to determine antibiotic resistance or susceptibility. These observations were further confirmed by the cell morphology (Figure 4C). Susceptible bacterial cells have a rounded shape, due to the difficulties with the outer wall synthesis which often resulted in the cell lysis, while resistant cells maintain a typical rod shape morphology and growth in micro-colonies or as planktonic cells. Images showing the morphology changes for one selected chamber per each tested antibiotic are in Figure S10.

Further, we tested the presented microfluidic platform using a clinical isolate of *E.coli*. We performed an AST for this isolate against meropenem, ciprofloxacin and amoxicillin at the concentrations identical to the previous experiments. The resulting resistance profiles are presented in Figure 5.

**Figure 5.**
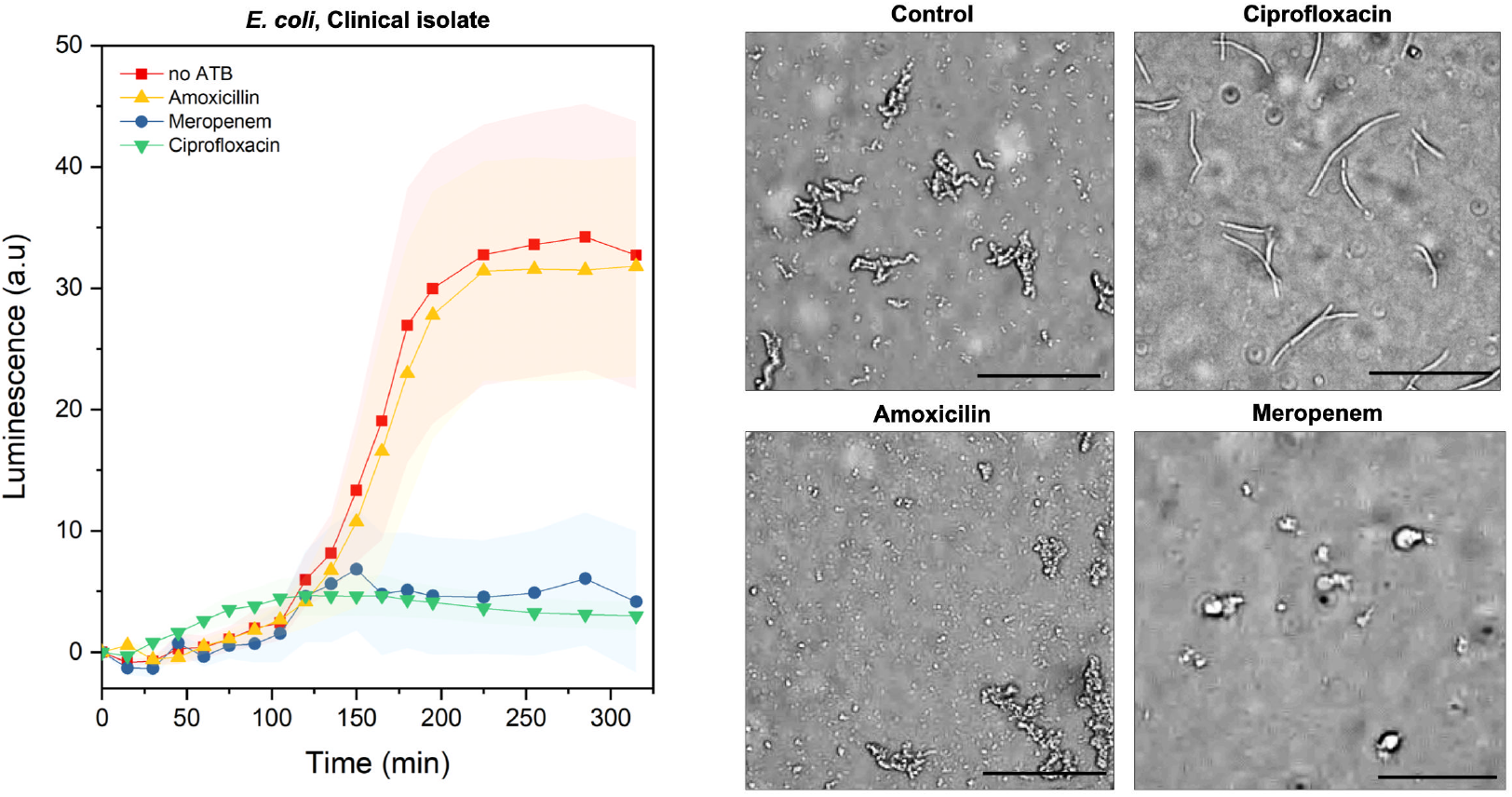
Validation of the AST platform using a clinical isolate. Kinetic cell viability measurements via oxygen sensing nanoprobes of *E. coli* clinical isolate. Comparison between the standard bacterial culture without antibiotics (RED, n=29) and cultivation in the presence of amoxicillin (YELLOW, n=48), meropenem (BLUE, n=33) and ciprofloxacin (GREEN, n=47) at the concentrations four times greater than reference MIC_ref_ for *E.coli*. Corresponding brightfield micrographs show the bacterial morphology at the end of the assay, after 5 hours of incubation. Inoculation was performed at OD_600_ of 0.02 (20 μL). Temperature: 37 °C, MHB II, cation adjusted. Error bars represent the standard deviations. Scale bars: 50 μm.

In comparison to the oxygen consumption for the culture supplied with an excess of different antibiotics and the control without the antibiotics, we could detect a reduced metabolic activity in the cultures containing meropenem and ciprofloxacin. We did not observe any decrease in the bacterial respiration in the culture containing amoxicillin. In contrast, the luminescence profile is similar to that of the culture without addition of the antibiotics, suggesting the resistance of this isolate against beta-lactam antibiotics. Additionally, we were able to confirm the obtained results by observing the cell morphology. In the case of meropenem we observed cell wall deformations; in case of ciprofloxacin we observed cell elongation.

In the culture exposed to amoxicillin, we observed cell growth, division and morphologies consistent with the control experiment, further confirming the resistance of the studied clinical isolate against amoxicillin. Altogether, based on the results of the AST performed on the microfluidic platform, the tested strain was susceptible to the ciprofloxacin and meropenem while resistant to the amoxicillin. These results were consistent with results we obtained by independent studies using the standard broth dilution technique (Table S1). The time-lapse images showing the morphology changes for one selected chamber per each tested condition can be found in the supplementary information (Figures S11, S12, S13 and S14).

## CONCLUSIONS

In conclusion, we presented simple to use microfluidic AST platform based on the luminescence mediated oxygen consumption assay, which was enabled through the use of the air-tight polymer COC. The response of the microbial cells was visible in real-time, showing the clear difference between the susceptible and the resistant bacterial strains.

We believe that preparation and filing of the chambers with gel, antibiotics and nanoprobes can be further optimized to be compatible with mass production and long-term storage. Once the chambers are prepared, they serve as miniaturized agar plates, ready for the inoculation. The bacterial cells are deposited to the chambers in a single step for the entire platform by simply pipetting culture on a glass slide, making the AST simple to perform by medical professionals without any further training. The required cell number is in a similar range as required for the state-of-the-art technologies, while we obtain a significantly higher number of technical replicates in parallel. We envision that in the future, we can incorporate even more antibiotics and their dilutions to the system with an identical footprint. The platform requires approximately 3 (repetitions) × 50 CFUs per condition; therefore, it is presumably possible to incorporate up to 48 different antibiotics at one platform and avoid long pre-culture times by filtering and hence up-concentrating the pre-culture. Additionally, the results can be obtained in about 2-3 hours of cultivation, while many state-of-an-art technologies and cultivation on agar plates require 16-24 hours to determine resistance profiles.

Here we used an automated microscope to visualize the cells and confirm the results of the nanoprobe-based signal during the system characterization, but the optical setup for the routine AST could be much more straightforward. The luminescence levels could be easily determined by implementing an optical fiber-based detector or coupling with a very simple optical setup. The recent advancement in the oxygen sensing nanoparticles^43^ could further increase the sensitivity of the oxygen-based sensing platforms and allow to simplify the optical setup for the luminescence quantification.

Potentially, the system could be used with different types of gel matrices, providing a possibility to identify and cultivate fastidious organisms and slow-growing pathogens. Similarly, other viability reagents could be incorporated to the gel matrix to allow AST for the anaerobic pathogens as well. Furthermore, the clamping device can be opened after initial assessment of cells and may allow further investigation of cells that were classified as “resistant”, hence opening the ways to investigate the underlying mechanisms of emergence of the resistance.

## MATERIALS AND METHODS

### Antibiotics for the AST assay

Meropenem trihydrate, ciprofloxacin hydrochloride monohydrate (stock solutions prepared in DI water) and amoxicillin (stock solution prepared in phosphate buffer, pH 6) were purchased from Sigma-Aldrich. Solutions of antibiotics in media were prepared shortly before the experiments, from aliquots of stock solutions (~500 μL) stored at the −20 °C. Each aliquot of the stock solutions was thawed only once, without refreezing.

### Antibiotics for plasmid maintenance

Kanamycin sulfate was obtained from the Sigma-Aldrich and stored in 1 mL aliquots (−20 °C). Ampicillin sodium salt was obtained from Glentham Life Sciences, dissolved in water and stored in 1 mL aliquots at the −20 °C.

### Bacterial strains

*Escherichia coli*, ATCC 25922 and ATCC 35218

These strains were modified to overexpress the gene for the superfolder green fluorescent protein, sfGFP, providing an additional means of monitoring cell viability and morphology changes induced by the presence of the antibiotic compounds. We transformed these strains with plasmid pSEVA271-*sfgfp* (lab collection) carrying the *sfgfp* gene, which was constructed based on plasmid pSEVA271^44^ (kanamycin resistance) adding a gene for sfGFP^45^ under control of a constitutive promoter (BioBrick part BBa_J23100).^46^

*Escherichia coli*, Clinical isolate

The strain used was obtained from the Division of Clinical Bacteriology and Mycology of the University Hospital Basel from routine diagnostics. The isolate originated from a positive blood culture. Blood culture samples were incubated with the Virtuo (bioMérieux) in respective blood culture bottles (BactAlert FA/FN, bioMérieux) for a maximum of 6 days. The positive blood culture was subcultured and the species were identified using MALDI-TOF mass spectrometry (microflex, Bruker).

### Fabrication of the COC chamber array plate

The fabrication process started with transferring the array design on to the initial master wafer. We used a standard SU-8 photolithography processes according to the manufacturer guidelines. In short, SU-8 3050 was spin coated at 2750 rpm for 30 s on a 4 inch silicon wafer substrate and subsequently soft baked resulting in a structure height of 75 μm. After exposure with UV light through a foil mask, the wafer was post-exposure baked and developed in a developer bath. Finally, the wafer was hard baked and silanized with trichloro (1H,1H,2H,2H-perfluorooctyl) silane (PFOTS) to avoid unwanted adhesion.

The stamp for the thermal imprinting was prepared by transferring the design from the initial master wafer to an UV curable resist (Ormostamp, Micro Resist, Germany) deposited on a 4 inch glass wafer. The resulting stamp was silanized with PFOTS. The nano-wells were thermally imprinted to the cyclic olefin copolymer (COC grade 8007) foil (Topas Advanced Polymers, Germany) with a thickness of 240 μm, which was cut to fit a 4 inch wafer format. The imprinting process was performed using a compact nanoimprinting tool (CNI V2.0, NILT, Denmark) by applying a pressure of 6 bar for 3 min at 130 °C. Finally, the COC wafers were diced into individual chips comprising four separate arrays each.

### Bacterial cultivation

Bacterial strains were stored as cryo-stocks containing 25% glycerol (−80 °C). Before the experiments, a small portion of the stock culture (10-20 μl) was transferred into 2 mL MHB II, cation-adjusted, containing, if necessary, appropriate antibiotics for plasmid maintenance (ATCC 25922 [pSEVA271-*sfgfp*]: 50 μg mL^-1^ kanamycin sulfate, ATCC 35218 [pSEVA271-*sfgfp*]: 50 μg mL^-1^ kanamycin sulfate and 100 μg mL^-1^ ampicillin sodium salt). The liquid culture was grown at 37 °C using a shaking incubator (Minitron, Infors HT) with a shaking speed of 220 rpm. When an OD_600_ of 0.2-0.4 was reached, the culture was diluted to OD_600_: 0.02 in the fresh cultivation medium.

The cultivation of the clinical isolate was performed under the similar conditions, within a biosafety level 2 laboratory (no antibiotics were used for the pre-cultivation).

### Clamping device

The layout of the clamping device was designed to fit a standard microscopy holder for a 96 well plate. The system (Figure S3) was composed from the two parts; the bottom part which closely fit the microscopy glass slide (24 x 40 mm, #5; Thermo scientific) and contained grooves to guide positioning of the top part precisely above the microfluidic device. Both parts had a set of the eight openings to fit block magnets (four 20 × 5 × 2 mm; four 10 × 3 × 2 mm; eight 6 × 4 × 2 mm, obtained from supermagnete.ch) used to hold both parts together. The top part had an opening in which a PDMS slab with a polymethyl methacrylate sheet on top was placed. The PDMS allowed an even distribution of the force exerted on the microfluidic platform by the magnets and keep it in close contact with the glass slide during incubation and measurement. Both parts were designed using the SOLIDWORKS 2019 software (Dassault systèmes) and printed using the Ultimaker 3 (Ultimaker) from ABS plastic.

### Preparation and deposition of the gel

The gel preparation process started with heating a 1.5 ml centrifugation tube (Eppendorf) with 100 μl aliquot of the ultra-low gelling temperature agarose gel solution (6% in Mueller Hinton Broth II, cation-adjusted) to 85 °C in a Thermomixer (Eppendorf Thermomixer R Shaker, Eppendorf). Once the temperature was reached, 20 μl of the OXNANO oxygen sensing nanoprobes (Pyroscience) was added to the agarose gel. Stock solution of the oxygen sensing nanoprobes was prepared by dissolving probes in DI water to a final concentration 1 mg mL^-1^; prior to the experiments, the stock solution was homogenized using the ultrasonic bath for at least 20-60 min.

The mixture of the gel and particles was heated using the thermomixer to 95 °C and held at the temperature for at least 5 min (to avoid potential bacterial contamination from the particle solution) and brought down to 47 °C. The solution of antibiotics in cultivation medium (2× the target concentration-viz table 1) was heated to 47 °C and 120 μl of the solution was added to the gel and mixed using a vortex mixer.

The solution of gel, particles and antibiotics was maintained at 47 °C; about 20 μl of the solution was pipetted over each array and spread using a small piece of PDMS. Once prepared, arrays were directly used for measurements.

During the process, all the chemicals were prepared and manipulation was performed in a sterile environment. Similarly, all the tools were sterilized before use to avoid contamination.

#### Modifications in the protocol

For the characterization of the system we used bacterial strains (ATCC 25922, ATCC 35218) constitutively producing the fluorescent protein sfGFP. To maintain its production, the growth medium additionally contained kanamycin sulfate (during the pre-culture and AST). The pre-culture of the ATCC 35218 *E. coli* strain contained ampicillin sodium salt in order to maintain the resistance. However, ampicillin was not supplied during the AST.

### Imaging of the microfluidic device

The inoculated microfluidic array was placed in the magnetic clamping system, which was directly placed on the holder of an automated, fully motorized inverted wide field microscope (Nikon Ti-E, controlled with NIKON NIS-Elements Advanced Research software) to perform time lapse imaging. The environmental box of the microscope was pre-heated to 37 °C and maintained at this temperature during the five-hour on-chip cultivation. We imaged individual chambers (chambers with entrapped air bubbles or visible damage of the gel were not imaged or evaluated) every 15 min for three hours and then for the following two hours every 30 min using brightfield and luminescence microscopy with use of the Nikon Perfect Focus System. Imaging was performed using a 10× objective (Nikon, Plan Fluor NA:0.3, WD:16 mm) with a Lumencor Spectra X LED light source for luminescence excitation using the following optical configurations: sfGFP fluorescence: cyan LED (30%), 475/28 excitation filter, 495 dichroic, 525/50 emission filter, exposure time 100 ms; oxygen sensor: blue LED (30%), 438/24 excitation filter, 660 dichroic, 785/60 emission filter, exposure time 200 ms. Images were recorded by a Hamamatsu Orca Flash 4 camera.

### Data analysis

The luminescence intensities from the individual chambers were analyzed using FIJI image analysis software.^47^ Regions of interest (ROI) (315 × 315 μm) were drawn around each chamber and the mean grey value was obtained for each of the 18 time points. Considering that bacterial cells as well nanoprobes were present also outside of the chambers, we did not perform a background subtraction, but instead subtracted the initial mean luminescence value obtained for the individual chambers at time point zero.

### AST *via* broth microdilution

The AST results from the microfluidic arrays were compared to the results obtained from a standard broth microdilution AST. This assay was performed following CLSI guidelines and as described previously^48^ but using a 384-well plate (40 μL final volume per well) instead of a 96-well plate format. The inoculum was prepared as described for the microfluidic assay but using a final cell concentration of 5 x 10^5^ CFU mL^-1^. The highest concentration of antibiotics was 800 μg mL^-1^ (amoxicillin), 1.6 μg mL^-1^ (meropenem) and 0.8 μg mL^-1^ (ciprofloxacin) in the first well which was then used for a 10-step serial dilution (2log). After incubation (37 °C, 18 h), the OD_600_ values of the plate were recorded using an Infinite 200 PRO plate reader (Tecan).

### Oxygen sensing control experiment (Figure S1)

To show the dependence of the luminescence signal on the oxygen level in the samples, we prepare two DI water samples with a different dissolved oxygen levels. One sample was saturated with the nitrogen gas for more than one hour and was expected to have a lower oxygen level compared to a DI water sample which was placed on a shaker (190 rpm, 24 °C). During the experiment, about 330 μL of the both samples were pipetted in the standard 96 well-plate (Thermo Fisher Scientific-NunclonTM Delta Surface, 96 flat bottom transparent polystyrene plates) together with the 30 μL of the stock 1mg mL^-1^ oxygen nanoprobes, immediately covered with the polyester sealing film (Starlab, Switzerland) and placed on the automated microscope to record the luminescence. The data was recorded using a 4× objective, 200 ms exposure time and 50% LED intensity. (All other experimental details as above in section: Imaging of the microfluidic device). The data collection started as the tape was removed, thus reinstating the oxygen level of both samples. We recorded luminesce levels every five minutes, from four wells and from two positions in each well. We used FIJI software to evaluate ROIs, square with edge size 1 mm. Measurements were performed at the room temperature, ~23 °C.

Measurements of the emission spectrum were performed on a Microplate Reader, Infinite M1000 PRO (Tecan) using Thermo Fisher Scientific-NunclonTM Delta Surface, 96 flat bottom transparent polystyrene plates. The sample with a lower oxygen level was prepared again by saturation with nitrogen gas and sealing immediately after preparing three wells with 360 μL sample containing 30 μl of the particle stock solution. The emission spectrum of these samples were compared to the samples without the pretreatment. The spectra for both samples between 500 to 850 nm were collected at an excitation wavelength of 438 nm in 5 nm steps. The chambers were sealed during the measurements. The spectra were recorded with three repetitions for each condition. Measurements were performed at ~30 °C.

## Author Contributions

The manuscript was written through contributions of all authors. All authors have given approval to the final version of the manuscript.

## Funding Sources

The financial support of the Swiss National Foundation SNF NRP72, project 407240_167123, and the European Research Council (ERC Consolidator grant no. 681587) are gratefully acknowledged. DGR is grateful for financial support from the Banting Postdoctoral Fellowship program.

## ACKNOWLEDGMENTS

The authors thank Tania Roberts (D-BSSE, ETH Zurich) for her valuable help with setting up experiments with the clinical isolate. We thank the Clean Room Facility and the Single-Cell Facility (both at D-BSSE, ETH Zurich) for their valuable help and advice, particularly Erica Montani, Tom Lummen and Mathias Markus Welsche.

## ABBREVIATIONS

ATB: antibiotic;
AST: antibiotic susceptibility testing;
COC: cyclic olefin copolymer;
MHB II: Mueller Hinton Broth II, cation adjusted;
MIC: minimal inhibitory concentration;
PDMS: polydimethylsiloxane;
PFOTS: trichloro (1H,1H,2H,2H-perfluorooctyl) silane ROI, region of interest;
sfGFP: superfolder variant of the green fluorescent protein

## Notes

### Competing Interest Statement

The authors have declared no competing interest.

