## Supporting information for "Real-time respiration changes as a viability indicator for rapid antibiotic susceptibility testing in a microfluidic chamber array"

### Supporting figures

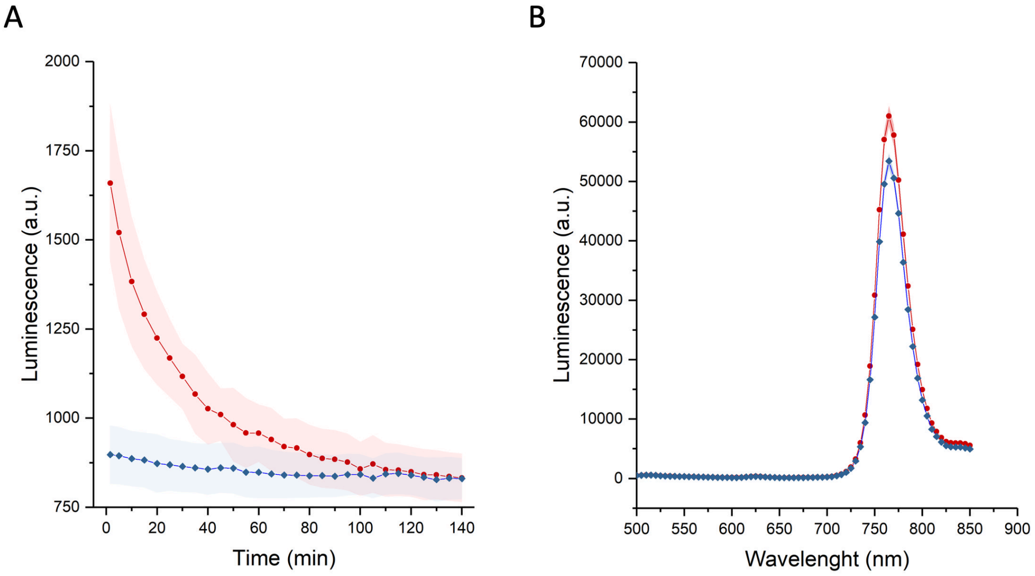


**Figure S1.** Comparison between two DI water samples with different dissolved oxygen levels containing oxygen sensing nanoprobes at a concentration identical to the AST measurements (83 µg $mL^{-1}$).
A: Time-resolved changes of the luminescence of the oxygen sensing nanoprobes measured on the 96 well-plate (n=8 for each sample) using an automated microscope.
The water sample saturated with nitrogen (red) shows a higher luminesce level compared against the sample without the pre-treatment (blue). All the chambers were open during the measurement; therefore, the oxygen level (luminescence) changed with time and in about two hours reached the same level in all of the observed chambers. Error bars represent the standard deviations.
B: Emission spectra (excitation 438 nm) of the water sample saturated with nitrogen (red) compared to the emission spectrum of the sample without the pre-treatment (blue). The sample saturated with the nitrogen reaches a greater luminesce level. n=3 for each condition. Error bars represent the standard deviations.

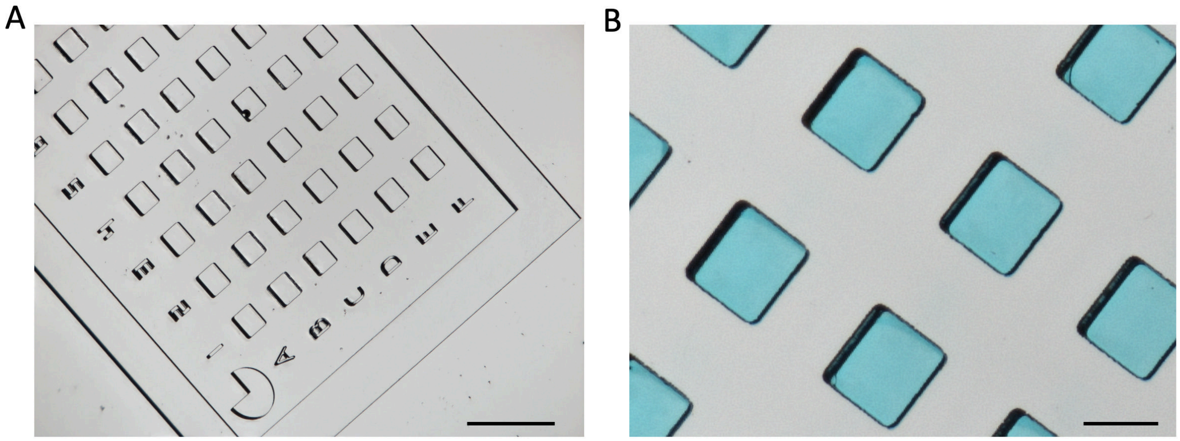


**Figure S2.** Microfluidic platform prepared in COC by thermal imprinting.
A: Detail figure of the one of the four arrays. Each row and column is marked with letters and numbers, respectively. Each of the four arrays also contains a symbol in the corner (here, 3/4 of the circle) allowing us to easily recognize each array (ATB concentration) during microscopy. B: Detail of the COC array filled with agarose gel containing blue food colorant for better visualization. Scale bars: 1mm, 250 µm respectively

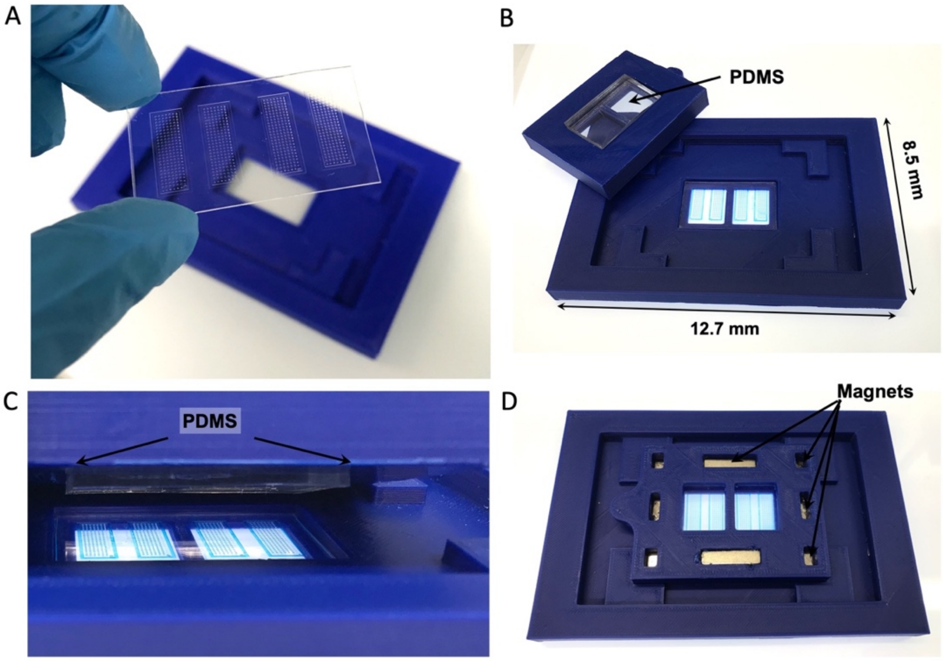


**Figure S3.** Clamping device: use and assembly.
The microfluidic array (A) is filled with agarose gel and placed over a sample-containing glass slide, positioned in the middle of the clamping device (B). The sample and the glass slide are kept in contact using the top part of the clamping device, which is pressing a PDMS slab (C) against the array using the magnetic force. The top and the bottom parts contain eight magnets in the corresponding positions, visible in picture D, showing the final assembled clamping system. (Agarose gel contains blue food colorant for visualization.)

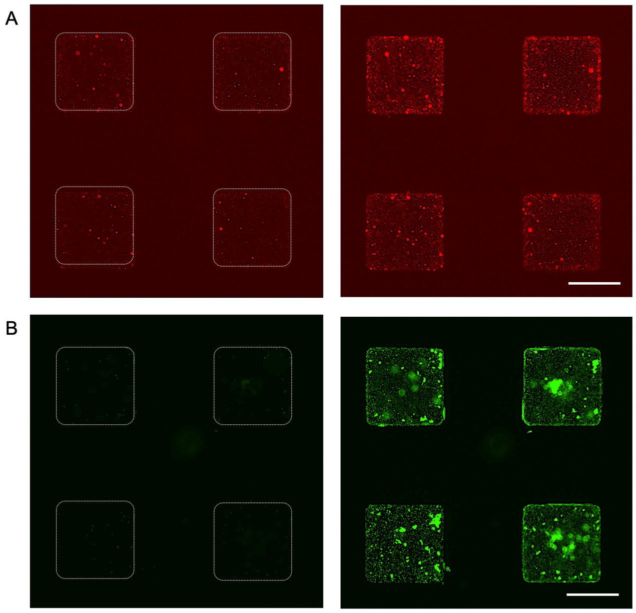


**Figure S4.** Four individual chambers of the AST microfluidic platform. (A) Detailed micrographs of the oxygen sensing nanoprobes embedded in the 2.5 % agarose gel at the beginning and at the end of the 5 h cultivation of the sfGFP producing *E. coli* ATCC 35218. (B) Detailed micrographs of the corresponding bacterial culture, again at the beginning and after the 5 h cultivation, visualized by the cell-produced sfGFP.
Scale bar: 200 µm

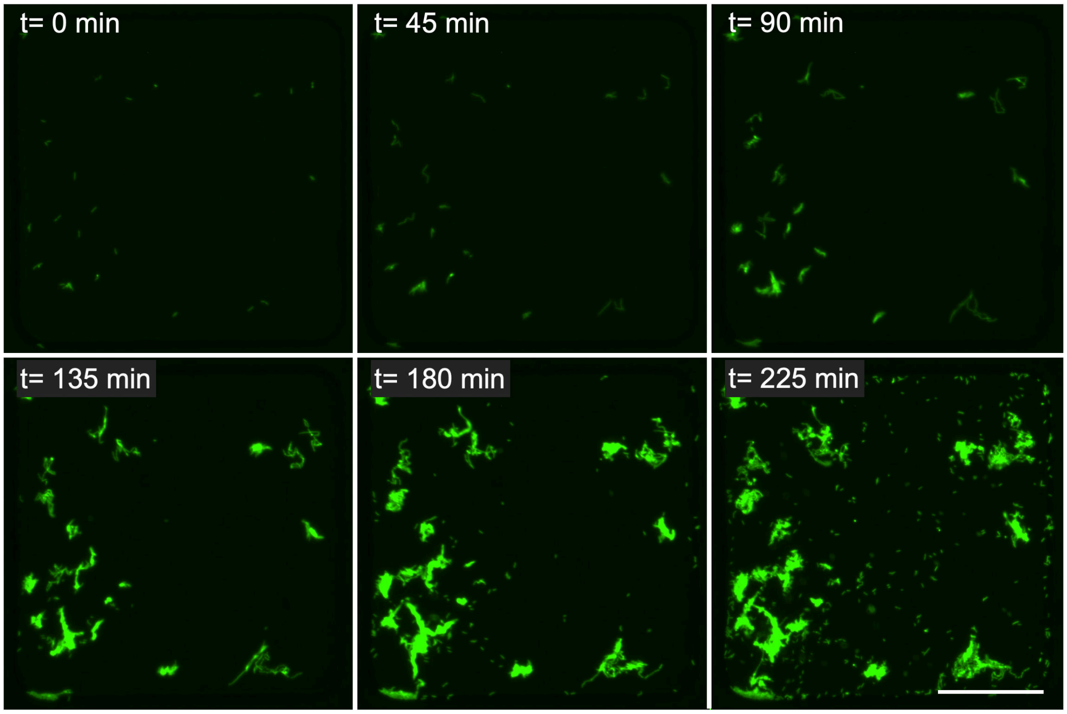


**Figure S5.** Time-lapse images from a selected chamber showing bacterial morphology (*E. coli* ATCC 25922 [pSEVA271-sfgfp]) during the cultivation in the microfluidic chamber system, here as a control without any antibiotics. Scale bar: 100 µm

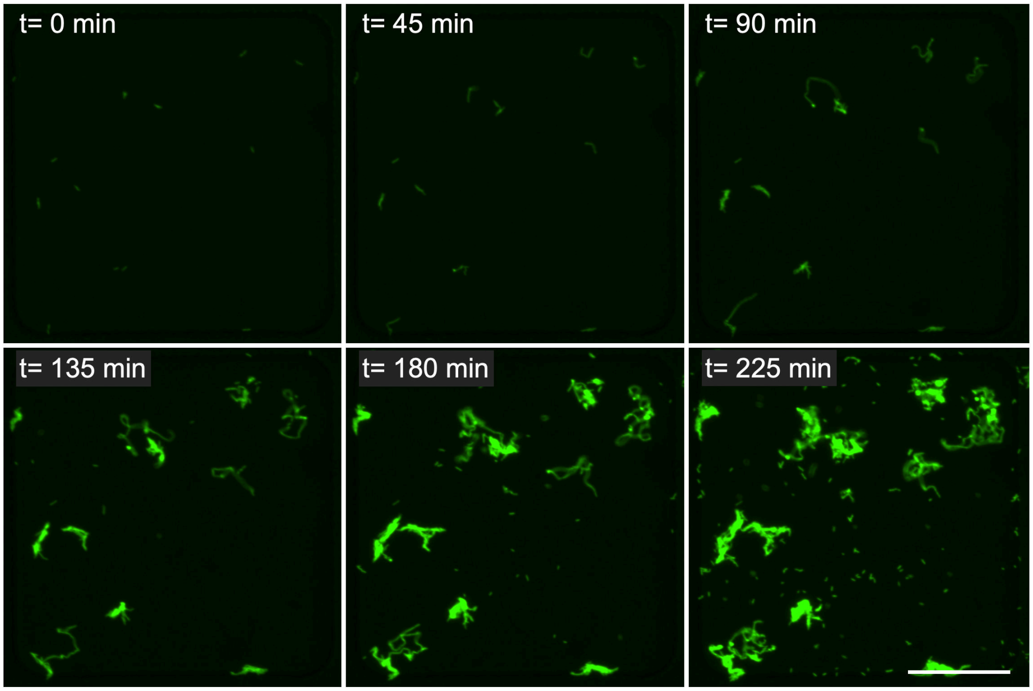


**Figure S6.** Time-lapse images from a selected chamber showing changes in bacteria morphology (*E. coli* ATCC 25922 [pSEVA271-sfgfp]) during cultivation in the microfluidic chamber system supplied with ciprofloxacin at a concentration of 0.25 $MIC_{ref}$. Measured from initial $OD_{600}$: 0.02; $MIC_{ref}$= 0.008 µg $mL^{-1}$. Scale bar: 100 µm

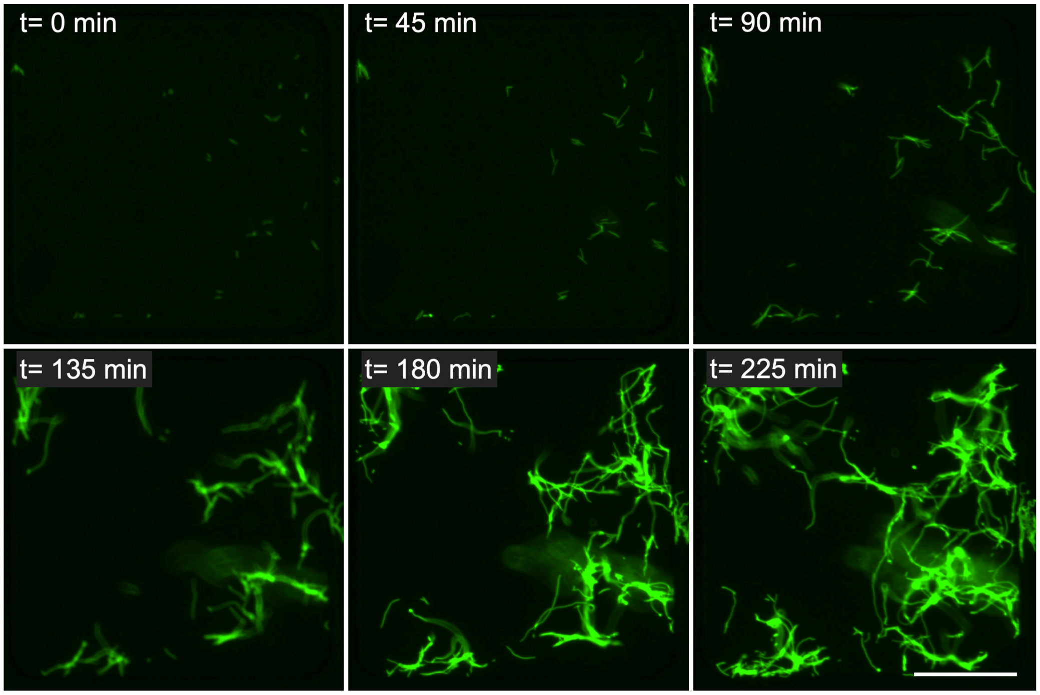


**Figure S7.** Time-lapse images from a selected chamber showing changes in bacterial morphology (*E. coli* ATCC 25922 [pSEVA271-sfgfp]) during cultivation using the presented microfluidic chamber system supplied with ciprofloxacin at a concentration of 1 $MIC_{ref}$. Measured from initial $OD_{600}$: 0.02; $MIC_{ref}$= 0.008 µg $mL^{-1}$. Scale bar: 100 µm

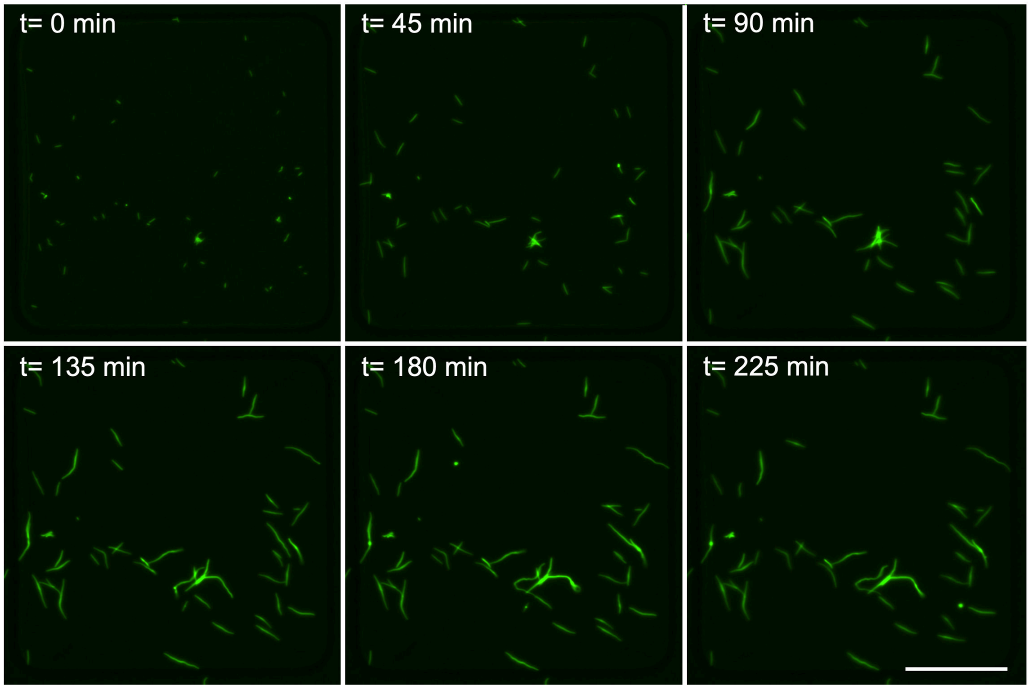


**Figure S8.** Time-lapse images from a selected chamber showing changes in bacterial morphology (*E. coli* ATCC 25922 [pSEVA271-sfgfp]) during cultivation using the presented microfluidic chamber system supplied with ciprofloxacin at a concentration of 4 $MIC_{ref}$. Measured from initial $OD_{600}$: 0.02; $MIC_{ref}$= 0.008 µg $mL^{-1}$. Scale bar: 100 µm

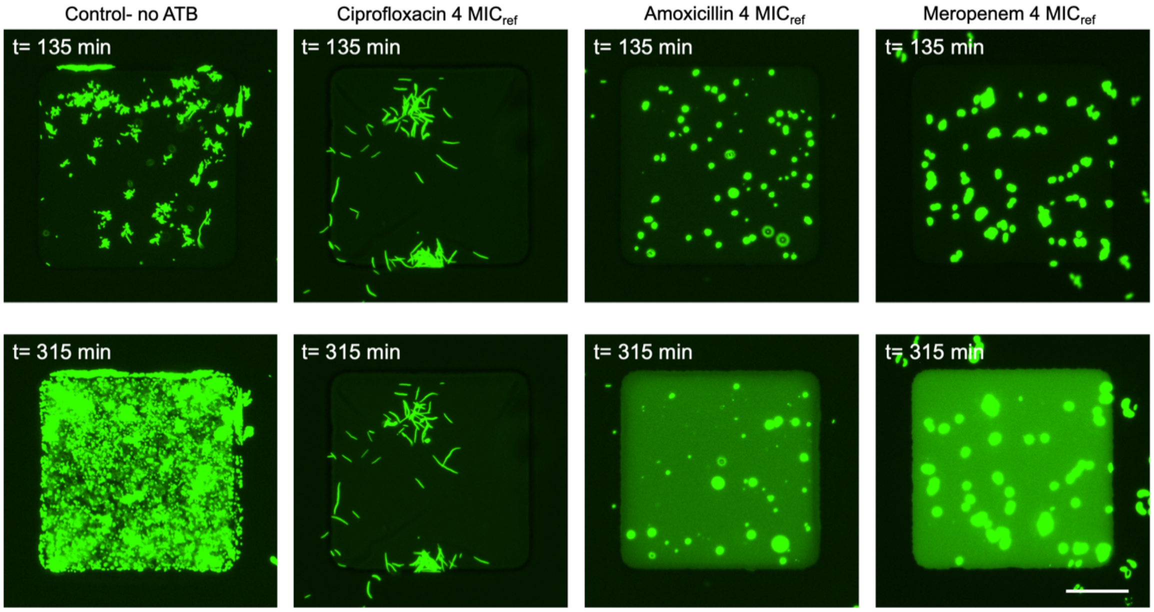


**Figure S9.** Images from selected chambers showing changes in bacterial morphology (*E. coli* ATCC 25922 [pSEVA271-sfgfp]) at two time points during cultivation using the presented microfluidic chamber system supplied with antibiotics at a concentration of 4 $MIC_{ref}$. Measured from initial $OD_{600}$: 0.02; ciprofloxacin-$MIC_{ref}$= 0.008 µg $mL^{-1}$, amoxicillin-$MIC_{ref}$= 4 µg $mL^{-1}$, meropenem-$MIC_{ref}$= 0.016 µg $mL^{-1}$. Scale bar: 100 µm

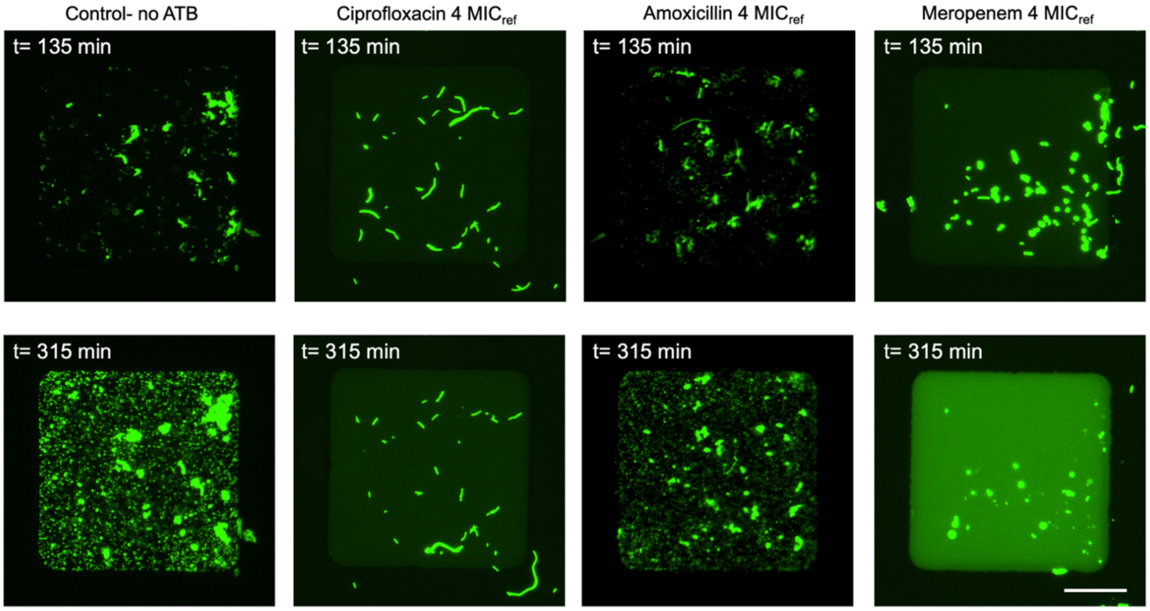


**Figure S10.** Images from selected chambers showing changes in bacterial morphology (*E. coli* ATCC 35218 [pSEVA271-sfgfp]) at the two time points of the cultivation using the presented microfluidic chamber system supplied with antibiotics at a concentration of 4 $MIC_{ref}$. Measured from initial $OD_{600}$: 0.02; ciprofloxacin-$MIC_{ref}$= 0.008 µg $mL^{-1}$, amoxicillin-$MIC_{ref}$= 4 µg $mL^{-1}$, meropenem-$MIC_{ref}$= 0.016 µg $mL^{-1}$. Scale bar: 100 µm

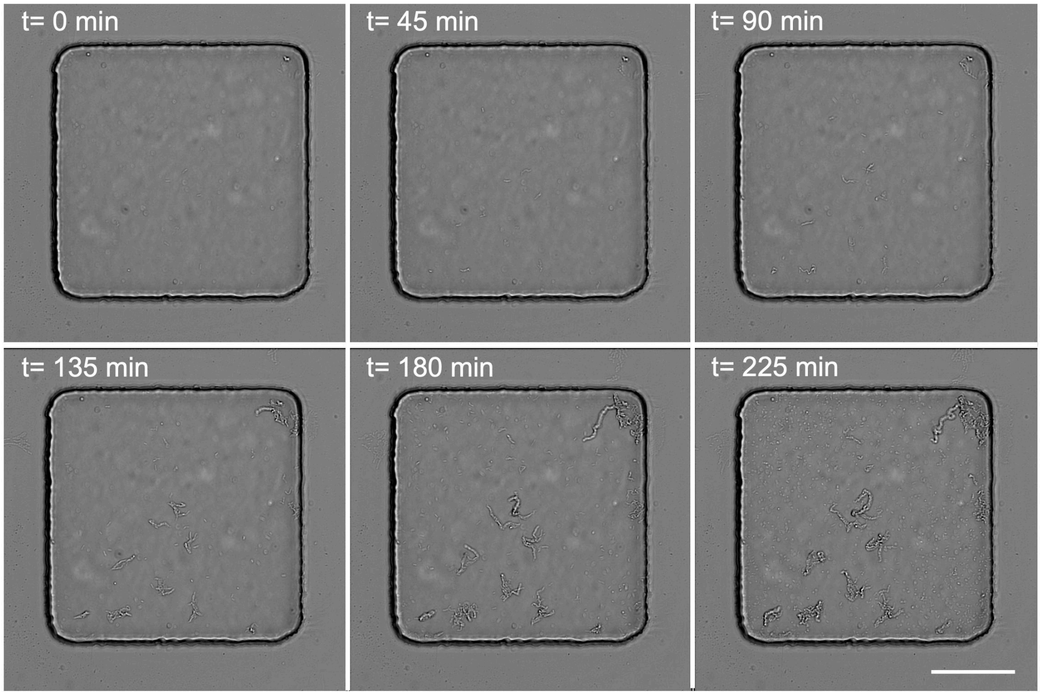


**Figure S11.** Time-lapse images from a selected chamber showing changes in bacterial morphology (*E. coli*-clinical isolate) during the cultivation without the antibiotic supply. Measured from initial $OD_{600}$: 0.02; Scale bar: 100 µm

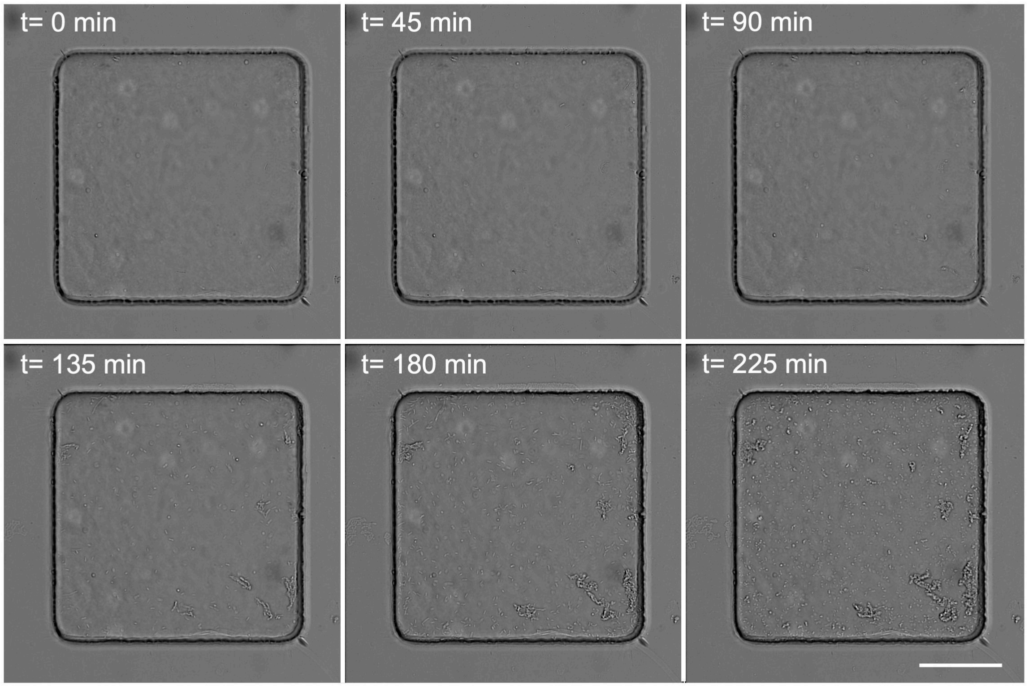


**Figure S12.** Time-lapse images from a selected chamber showing changes in bacterial morphology (*E. coli*-clinical isolate) during the cultivation in the presence of amoxicillin at the concentration 4 $MIC_{ref}$. ($MIC_{ref}$= 4 µg $mL^{-1}$). Measured from initial $OD_{600}$: 0.02; Scale bar: 100 µm

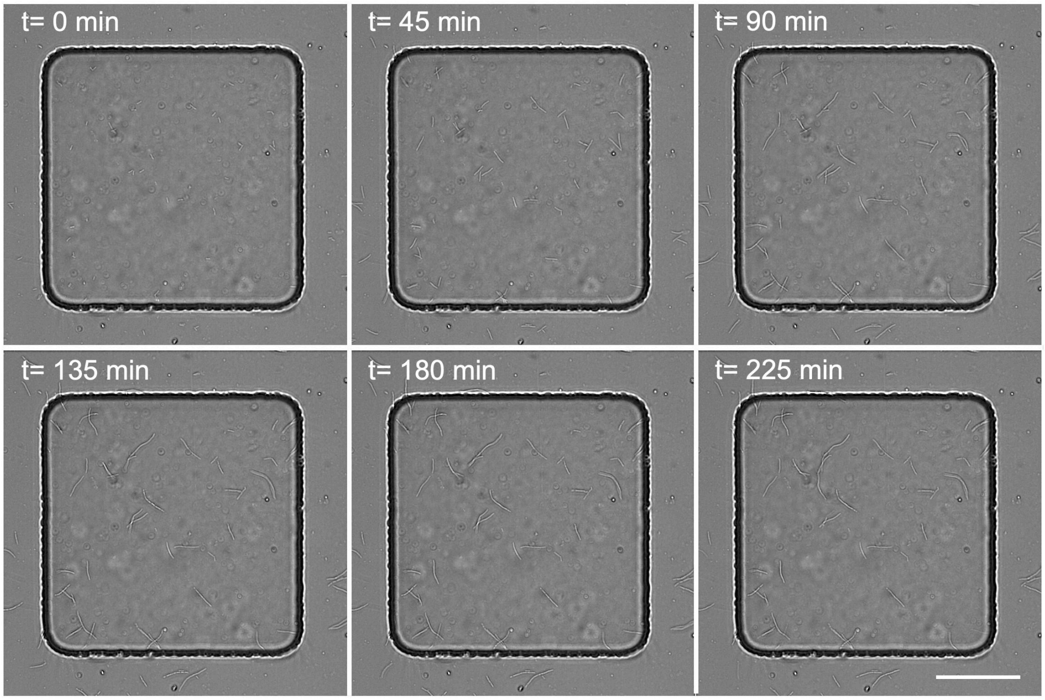


**Figure S13.** Time-lapse images from a selected chamber showing changes in bacterial morphology (*E. coli*- clinical isolate) during the cultivation in the presence of ciprofloxacin at the concentration 4 $MIC_{ref}$. ($MIC_{ref}$= 0.008 µg $mL^{-1}$). Measured from initial $OD_{600}$: 0.02; Scale bar: 100 µm

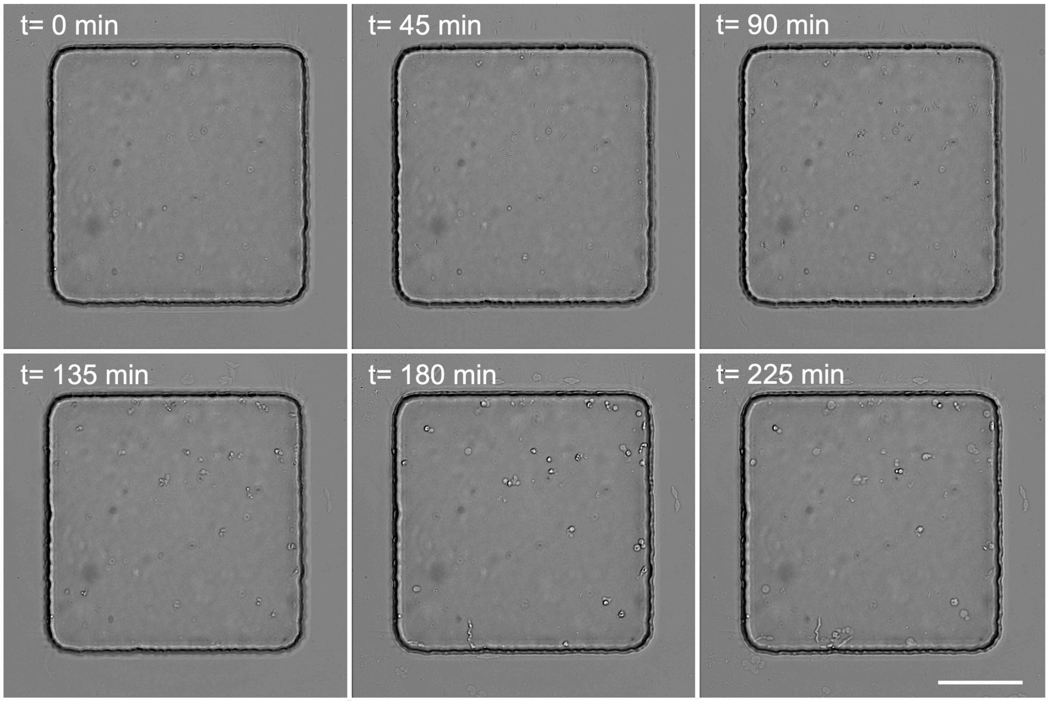


**Figure S14.** Time-lapse images from a selected chamber showing changes in bacterial morphology (*E. coli*- clinical isolate) during the cultivation in the presence of meropenem at the concentration 4 $MIC_{ref}$. ($MIC_{ref}$= 0.016 µg $mL^{-1}$). Measured from initial $OD_{600}$: 0.02; Scale bar: 100 µm

### Supplemental table

Table 1. MIC values for the studied *E. coli* strains.

| Bacterial strain | MIC Broth dilution [µg/ml] | | | MIC Broth/Agar dilution EUCAST [µg/ml] | | |
|---|---|---|---|---|---|---|
| | Amoxicillin | Meropenem | Ciprofloxacin | Amoxicillin | Meropenem | Ciprofloxacin |
| ATCC 25922 | 3.65 | 0.025 | 0.01 | 8.0/4.0 | 0.016 | 0.008 |
| ATCC 25922 [pSEVA271-sfGFP] | 4.17 | 0.025 | 0.008 | / | / | / |
| ATCC 35218 | / | 0.025 | 0.015 | / | / | / |
| Clinical isolate | Not active | 0.017 | 0.012 | / | / | / |
